# *DLG2–DLG4* Expression is Associated with Improved Survival and a Synaptic Gene Signature in Lower-Grade Glioma

**DOI:** 10.64898/2026.04.13.718176

**Authors:** Felipe Gaia, Henrique Ritter Dal-Pizzol, Osvaldo Malafaia, Rafael Roesler, Gustavo R. Isolan

## Abstract

**Background/Objectives:** Increasing evidence indicates that gliomas co-opt mechanisms of excitatory synaptic transmission and plasticity to support tumor progression, yet these processes remain poorly characterized in lower-grade gliomas (LGGs). Here, we investigated whether genes associated with excitatory synaptic function are linked to patient prognosis in LGG.

**Methods:** A curated panel of 36 synaptic genes was analyzed in LGG using RNA-sequencing and clinical data from The Cancer Genome Atlas (TCGA) and Chinese Glioma Genome Atlas (CGGA) datasets.

**Results:** Among the genes investigated, *DLG2, DLG3*, and *DLG4*, which encode the postsynaptic scaffolding proteins PSD-93, SAP-102, and PSD-95, respectively, showed strong associations with patient overall survival (OS). Higher expression of each gene was consistently associated with longer OS across both datasets. Expression of *DLG2*–*DLG4* was higher in oligodendroglioma and IDH-mutant, 1p/19q co-deleted tumors, and lower in astrocytoma and IDH-wild-type tumors. Furthermore, expression of all three genes positively correlated with a broad gene signature associated with a synaptic gene program, including multiple components of glutamatergic signaling and postsynaptic organization.

**Conclusions:** These findings suggest that elevated expression of *DLG2*–*DLG4* is associated with a transcriptional program resembling differentiated neuron-like features and favorable clinical outcome in LGG.

**Simple Summary:** Lower-grade gliomas are brain tumors with highly variable outcomes, and better markers are needed to predict how patients will fare. Recent research suggests that these tumors may use mechanisms normally involved in communication between brain cells, but this is not well understood in these cancer types. In this study, we analyzed large patient datasets to examine genes related to synaptic function. We found that higher expression of three genes involved in synaptic membrane organization, *DLG2, DLG3*, and *DLG4* was consistently associated with longer patient survival. These genes were also linked to a broader pattern of gene expression suggestive of neural transmission and plasticity. Our findings suggest that some lower-grade gliomas may adopt characteristics of normal brain cells that are associated with less aggressive behavior. This work may help guide future research on prognostic markers and improve understanding of brain tumor biology.

## 1. Introduction

Increasing evidence supports the view that tumor progression hijacks fundamental cellular mechanisms underlying neural transmission and synaptic plasticity. In particular, malignant gliomas exploit signaling pathways normally involved in excitatory synaptic communication, establishing functional interactions with surrounding neural circuits. In glioblastoma (GBM), the most aggressive primary brain tumor in adults, tumorigenesis, growth, and invasion are influenced by the formation of functional neuron–glioma synapses mediated by glutamate receptor-initiated signaling. Pioneering studies demonstrated that glioma cells receive direct synaptic input from neurons through α-amino-3-hydroxy-5-methyl-4-isoxazolepropionic acid (AMPA) glutamate receptors, leading to membrane depolarization and activity-dependent tumor proliferation. These neuron-to-glioma synapses enable electrical and synaptic integration of tumor cells into neural circuits, thereby coupling neuronal activity to tumor growth dynamics [1, 2].

In addition to simple synaptic coupling, glioma cells appear to recruit mechanisms of activity-dependent plasticity. Glioma synapses recruit canonical pathways of adaptive synaptic potentiation, supporting tumor progression through activity-driven processes [3]. These observations support a model in which glioma cells adopt and repurpose core components of synaptic transmission and plasticity to sustain tumor growth. This conceptual framework raises the possibility that genes classically associated with postsynaptic organization and excitatory synapse function may define biologically and clinically relevant programs in glioma.

Most studies showing parallels between gliomas and normal synapses focus on GBM, whereas synaptic mechanisms in lower grade glioma (LGG) types remain poorly understood. LGGs are generally less aggressive than primary GBM and occur in adults at an earlier age, with the highest incidence observed between the third and fourth decades of life. These neoplasms are characterized by relatively slow growth and infiltrative behavior, most commonly involving the cerebral hemispheres. Despite their initially indolent clinical course, LGGs frequently undergo progressive malignant transformation over time, contributing to significant neurological morbidity and mortality [4-6].

In the present study, we selected a panel of genes involved in excitatory synaptic transmission based on the available literature. We then analyzed whether these genes are associated with patient prognosis, assessed by survival, using datasets comprising GBM and LGG. Among the genes analyzed, *DLG2, DLG3*, and *DLG4* were among the ones showing particularly strong associations with survival specifically in LGG tumors, which were then chosen as the focus of our analyses. Interestingly, higher expression of each of these three *DLG* genes was associated with better, rather than worse, survival. *DLG2, DLG3*, and *DLG4* encode PSD-93 (SAP-93), SAP-102, and PSD-95, respectively, which are members of the membrane-associated guanylate kinase (MAGUK) family of synaptic scaffolding proteins. These proteins help organize the postsynaptic density through interactions with other scaffolding proteins and regulate the clustering of glutamate receptors [7]. PSD-93 is a key postsynaptic protein that links *N*-methyl-D-aspartate (NMDA) receptor activity to downstream signaling complexes and can form heterodimers with PSD-95, enhancing neural plasticity [8, 9]. PSD-95 also plays a central role in AMPA receptor trafficking during the induction of long-term potentiation (LTP) of excitatory synaptic transmission [10–12]. During early neurodevelopment, initial synapse formation is strongly influenced by SAP-102, whose role is later largely replaced by PSD-95 [7]. Following the initial screening, we then examined whether expression of *DLG2*–*DLG4* positively correlates with that of the other genes in our panel which are associated with synaptic transmission and plasticity.

## 2. Materials and Methods

### 2.1 Gene Panel for Excitatory Synaptic Transmission and Plasticity

Based on a curation of the available literature, we assembled a panel of 36 genes with key roles in excitatory synaptic transmission and synaptic plasticity (Supplementary Table S1).

### 2.2 Datasets and Gene Expression Analyses

RNA-sequencing and corresponding clinical data for patients with LGG were obtained from two independent cohorts that include World Health Organization (WHO) grade 2 and grade 3 gliomas, The Cancer Genome Atlas (TCGA-LGG, https://www.cancer.gov/tcga) [13] and the Chinese Glioma Genome Atlas (CGGA, http://www.cgga.org.cn) [14]. For the TCGA cohort, gene-level expected counts were retrieved from the TOIL hub using the UCSCXenaTools package, and clinical annotations were obtained from the TCGA Pan-Cancer Atlas dataset via cBioPortalData. For the CGGA cohort, both RNA-sequencing and corresponding clinical data from two datasets (CGGA-325 and CGGA-693) were downloaded and merged by retaining genes shared across both datasets and merged by retaining common genes. Batch (CGGA-325 vs. CGGA-693) was incorporated as a covariate in the DESeq2 design formula to account for technical variability during differential expression analysis. Samples with missing survival data or incomplete clinical annotations were excluded.

TCGA data were back-transformed from expected counts to ensure compatibility with count-based statistical modeling using DESeq2. The raw count matrices from the two CGGA datasets were merged after matching common genes. Duplicate gene symbols were resolved by retaining the entry with the highest mean expression across samples. Variance-stabilizing transformation was applied to normalized count data using DESeq2, and z-score scaling was subsequently performed on a gene-wise basis across all samples within each cohort.

Differences in the expression of *DLG2, DLG3*, and *DLG4* were evaluated across TCGA histological types [13, 15, 16], namely astrocytoma (TCGA, n = 188; CGGA, n = 239), oligodendroglioma (TCGA, n = 179; CGGA, n = 145), and oligoastrocytoma (TCGA, n = 123; CGGA, n = 23), and the updated classification incorporating molecular features (molecular subtypes) [17], namely LGG with a mutation in the isocitrate dehydrogenase (IDH) gene and co-deletion of chromosomal arms 1p and 19q (LGG-IDH-mut-codel; TCGA, n = 164; CGGA, n = 113), LGG with a mutation in the IDH gene without co-deletion of chromosomal arms 1p and 19q (LGG-IDH-mut-non-codel; TCGA, n = 235, CGGA, n = 142), and LGG wild-type for IDH (LGG-IDH-wt; TCGA, n = 91; CGGA, n = 88). Pairwise comparisons between groups were conducted using the non-parametric Wilcoxon rank-sum test, with *p*-values being adjusted for multiple testing via the Benjamini-Hochberg method.

For visualization of *DLG2-DLG4* expression in glioma compared to non-tumor brain tissue, the Rembrandt dataset [18] was analyzed using the GlioVis - Data Visualization Tools for Brain Tumor Datasets (https://gliovis.bioinfo.cnio.es/) platform [19].

### 2.3 Principal Component Analysis

Principal Component Analysis (PCA) was performed using the Evergene platform (https://bshihlab.shinyapps.io/evergene/) [20] to allow the visualization of *DLG2-DLG4* expression across the different histological types and molecular subtypes of LGG in relation to the first principal component (PC1), which captures the largest proportion of variance in the LGG transcriptomic data. Classification of TCGA LGG tumors within the Evergene platform is: histological types, astrocytoma, oligodendroglioma, oligoastrocytoma, or not available (NA); molecular subtypes, pilocytic astrocytoma (PA)–like, mesenchymal-like, glioma CpG island methylator phenotype (G-CIMP)-high, G-CIMP-low, glioma with 1p/19q co-deletion (codel), classic-like, or not available (NA).

### 2.4 Survival Analysis

Associations between overall survival (OS) of patients with LGG and expression analyses of the selected genes were assessed using Kaplan–Meier survival curves. Patients were dichotomized into “high” and “low” expression groups based on an optimal cutpoint determined by using the surv_cutpoint function in the survminer package (Supplementary Table S2) [12]. Survival distributions were compared using the log-rank test, with p-values adjusted for multiple testing across all genes and cohorts using the Benjamini–Hochberg method.

All analyses were performed in the R statistical environment (version 4.5.1). Variance-stabilizing transformation and batch adjustment were conducted using DESeq2 (version 1.48.1). OS analyses were performed using the survival (version 3.8.3), with optimal expression cutpoints determined using maximally selected rank statistics implemented in survminer (version 0.5.2). Kaplan–Meier curves were generated using survminer. Graphical visualizations were produced using ggplot2 (version 4.0.2). Data acquisition was performed using UCSCXenaTools (version 1.7.0), cBioPortalData (version 2.20.0).

### 2.5 Multivariate Cox Regression Analysis

Survival associations were evaluated using multivariable Cox proportional hazards regression models. Gene expression status for *DLG2, DLG3*, and *DLG4* was dichotomized into high- and low-expression groups as described. For each gene, Cox regression models were constructed including the following clinicomolecular covariates: IDH mutation status (mutant vs. wild-type), 1p/19q codeletion status (codel vs. non-codel), age included as a continuous covariate, WHO tumor grade (grade 2 or grade 3). Hazard ratios (HRs) and 95% confidence intervals (95% CIs) were estimated for each variable. Results were visualized using forest plots to illustrate the independent prognostic contribution of each co-variate within the multivariable model.

### 2.6 Gene Expression Correlations

Correlations between expression of each of the three selected *DLG* genes and each of the genes in the synaptic gene panel, or between each *DLG* gene and all genes in the panel pooled together to represent a synaptic signature, were analyzed with the Evergene platform using default settings [20]. Pearson correlation coefficient (*r*) values equal or higher than 0.5 along with *p*-values of less than 0.05 were considered to indicate moderate to high correlation.

For each gene in the synaptic signature, expression values were standardized across samples using z-score transformation. Pairwise gene–gene correlations were computed using Spearman’s rank correlation coefficient to capture monotonic relationships independent of distributional assumptions. A symmetric gene–gene correlation matrix was generated representing co-expression relationships among synaptic genes. For hierarchical clustering and visualization, the correlation matrix was hierarchically clustered using Ward’s minimum variance method. A heatmap was generated using the pheatmap package in R. Color scaling represented correlation strength ranging from negative (blue) to positive (red) values. No imputation was performed and missing values were handled by pairwise deletion.

## 3. Results

### 3.1 DLG2-DLG4 Expression Across Different LGG Types

An early analysis of associations of the genes in the synaptic panel with LGG patient survival revealed *DLG2, DLG3*, and *DLG4* among the genes with the most significant association. Their expression was then analyzed in LGG tumors classified into histological types or molecular subtypes. There was an apparent lower expression of *DLG2-DLG4* in glioma tumors compared to non-tumor brain tissue (Supplementary Figure S1). Within histological types, expression of *DLG2* in TCGA tumors was lower in astrocytoma compared to oligodendroglioma and oligoastrocytoma. *DLG3* and *DLG4* were more highly expressed in oligodendroglioma and less expressed in astrocytoma (Figure 1). Within the molecular classification, all three *DLG* genes showed the same pattern of higher expression in LGG-IDH-mut-codel and lower expression in LGG-IDH-wt, with intermediate levels in LGG-IDH-mut-non-codel (Figure 2).

**Figure 1.**
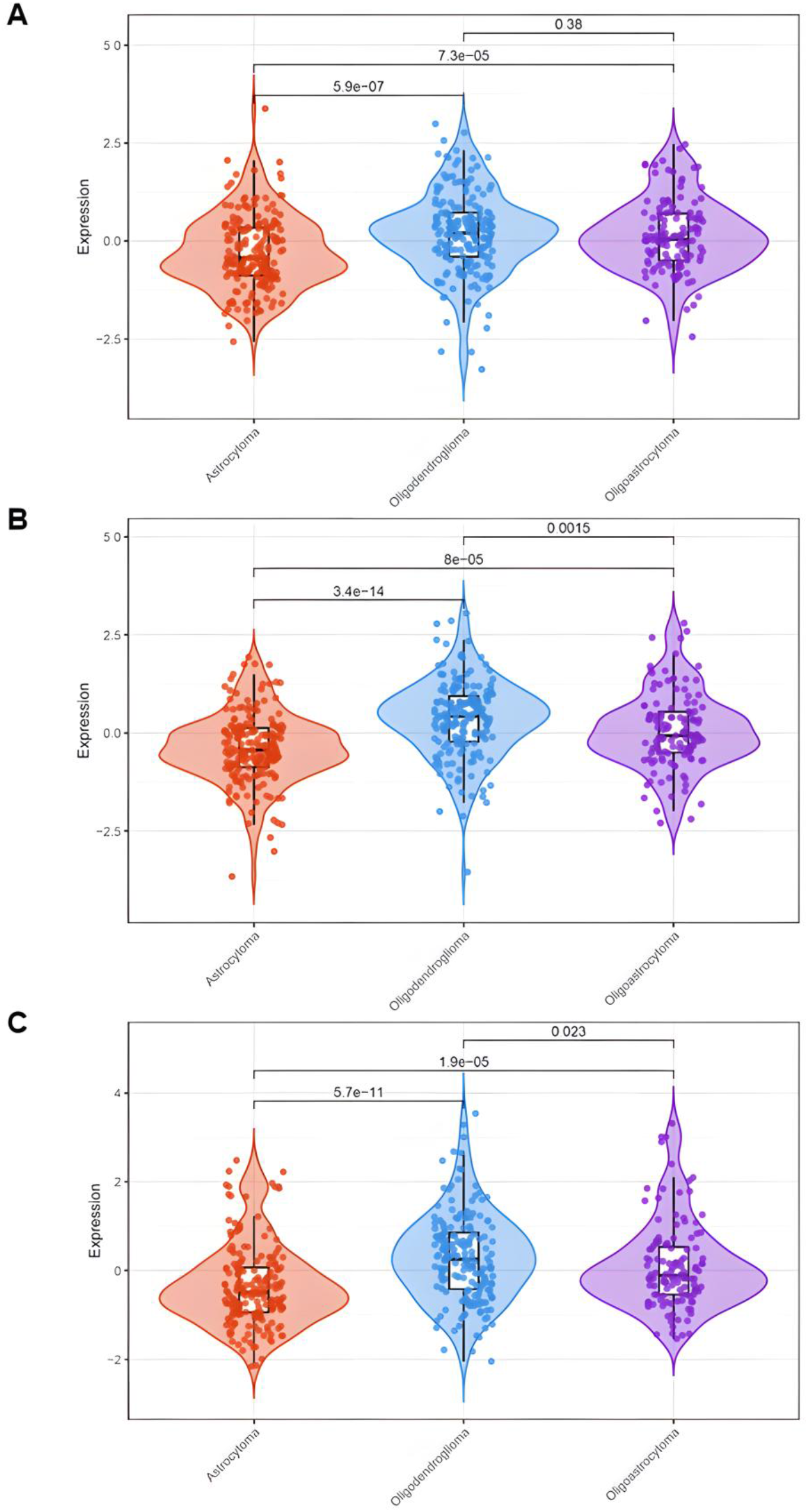
Gene expression levels of (**A**) *DLG2*, (**B**) *DLG3*, and (**C**) *DLG4* in TCGA LGG tumors classified into histological types. Astrocytoma, n = 188; oligodendroglioma, n = 179; oligoastrocytoma, n = 123; *p-*values are indicated in the panels.

**Figure 2.**
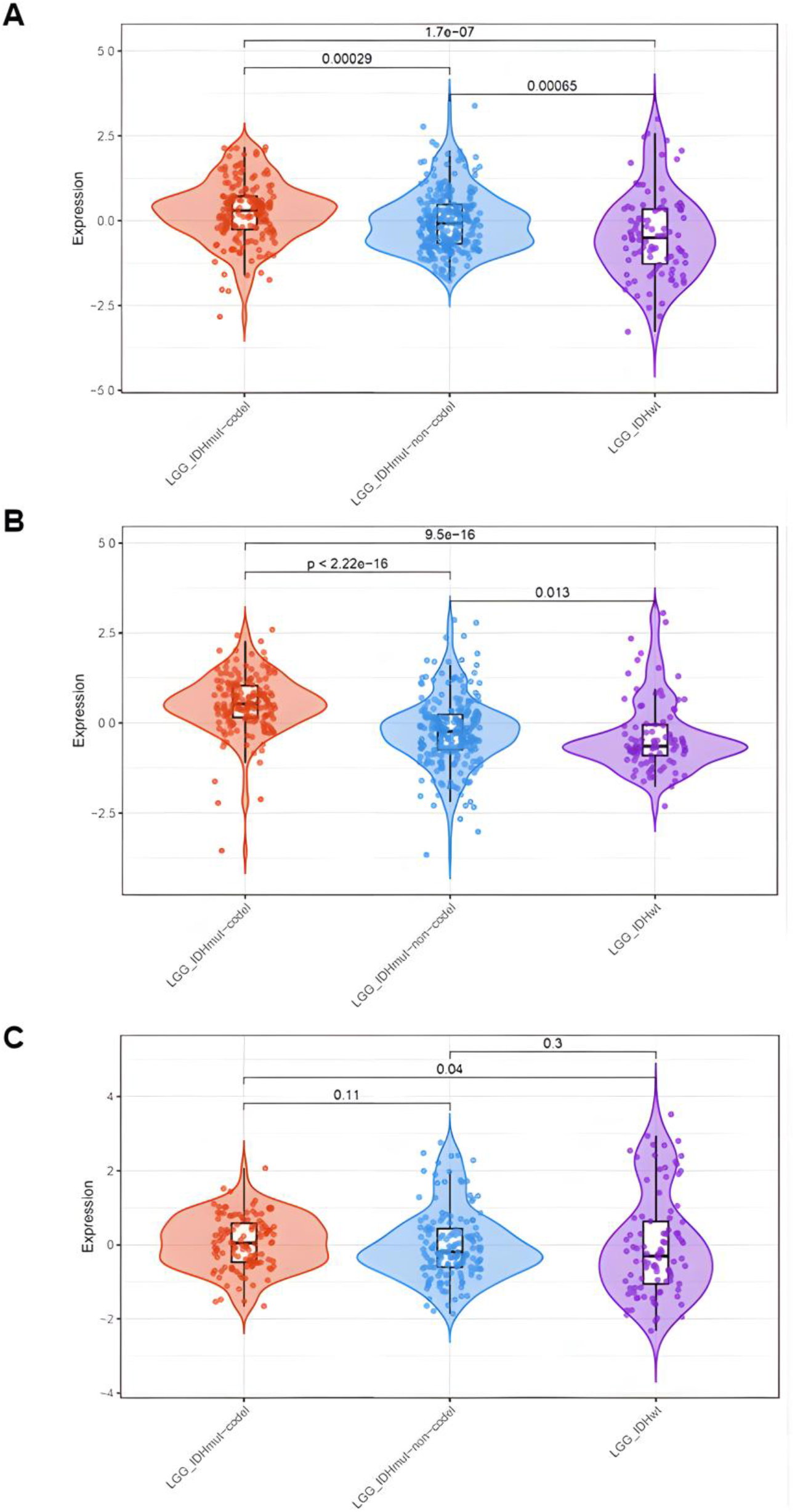
Gene expression levels of (**A**) *DLG2*, (**B**) *DLG3*, and (**C**) *DLG4* in TCGA LGG tumors classified into molecular subtypes. LGG-IDH-mut-codel, n = 164; LGG-IDH-mutnon-codel, n = 235; LGG-IDH-wt, n = 91; *p*-values are indicated in the panels.

Analysis of the CGGA dataset largely validated the overall pattern of lower *DLG2-DLG4* expression in astrocytoma compared to the other histological groups, and *DLG3* was more highly expressed in oligoastrocytoma than in oligodendroglioma (Supplementary Figure S2). In CGGA tumors classified according to molecular subtype, *DLG3* and *DLG4* were more expressed in LGG-IDH-mut-codel, but there were no statistically significant differences among tumor types in *DLG2* expression, although there was a trend towards higher expression in LGG-IDH-mut-codel compared to LGG-IDH-mut-non-codel (Supplementary Figure S3). The relationship between *DLG2*–*DLG4* expression and PC1 in TCGA tumors was illustrated by PCA, showing an overall pattern of greater alignment with PC1 in tumors with lower *DLG* expression levels (Supplementary Figure S4). In TCGA LGG tumors, the PC1 gene set (*BCL2L12, S100A11, TMSB4X, CLIC1, RAB32, RNF135, CASP1, SP100, AGTRAP, SQOR, CARD16, VIM, RHOC, KDELR1, AK2, CD58, ZDHHC12, RER1, SERTAD3, APOBEC3G*) is predominantly associated with a mesenchymal, inflammatory, and stress-response axis, related to inflammation, immune evasion, reactive glial phenotype, cell motility, cytoskeleton remodeling, resistance to apoptosis, and tumor–microenvironment interactions [13].

### 3.2 High DLG2-DLG4 Expression is Associated with Longer OS in Patients with LGG

High expression of *DLG2, DLG3*, and *DLG4* was significantly associated with longer OS in patients with LGG in both the TCGA (and CGGA datasets (Figure 3). Multivariable Cox proportional hazards analyses demonstrated that expression levels of *DLG2-DLG4* genes retained significant prognostic associations in the CGGA dataset, but not in the TCGA dataset, after adjustment for established clinicomolecular variables, including IDH mutation status, 1p/19q codeletion status, age, and tumor grade. Forest plot visualization showed the relative contribution of each covariate to overall survival risk. As expected, IDH mutation and 1p/19q codeletion were associated with more favorable outcomes, whereas increasing age and higher tumor grade were associated with poorer survival. High expression of *DLG2, DLG3*, and *DLG4* was associated with reduced hazard of death in CGGA tumors, supporting an independent favorable prognostic role for synaptic-associated *DLG* gene expression in LGG. However, although the *DLG* genes in TCGA tumors showed a similar trend, confidence intervals crossing the null hazard ratio, indicating lack of statistical significance within the multivariable model. Thus, *DLG2-DLG4* expression in TCGA tumors was associated with favorable survival in univariate analyses, but this association was attenuated after adjustment for established clinicomolecular variables (Supplementary Figure S5, S6).

**Figure 3.**
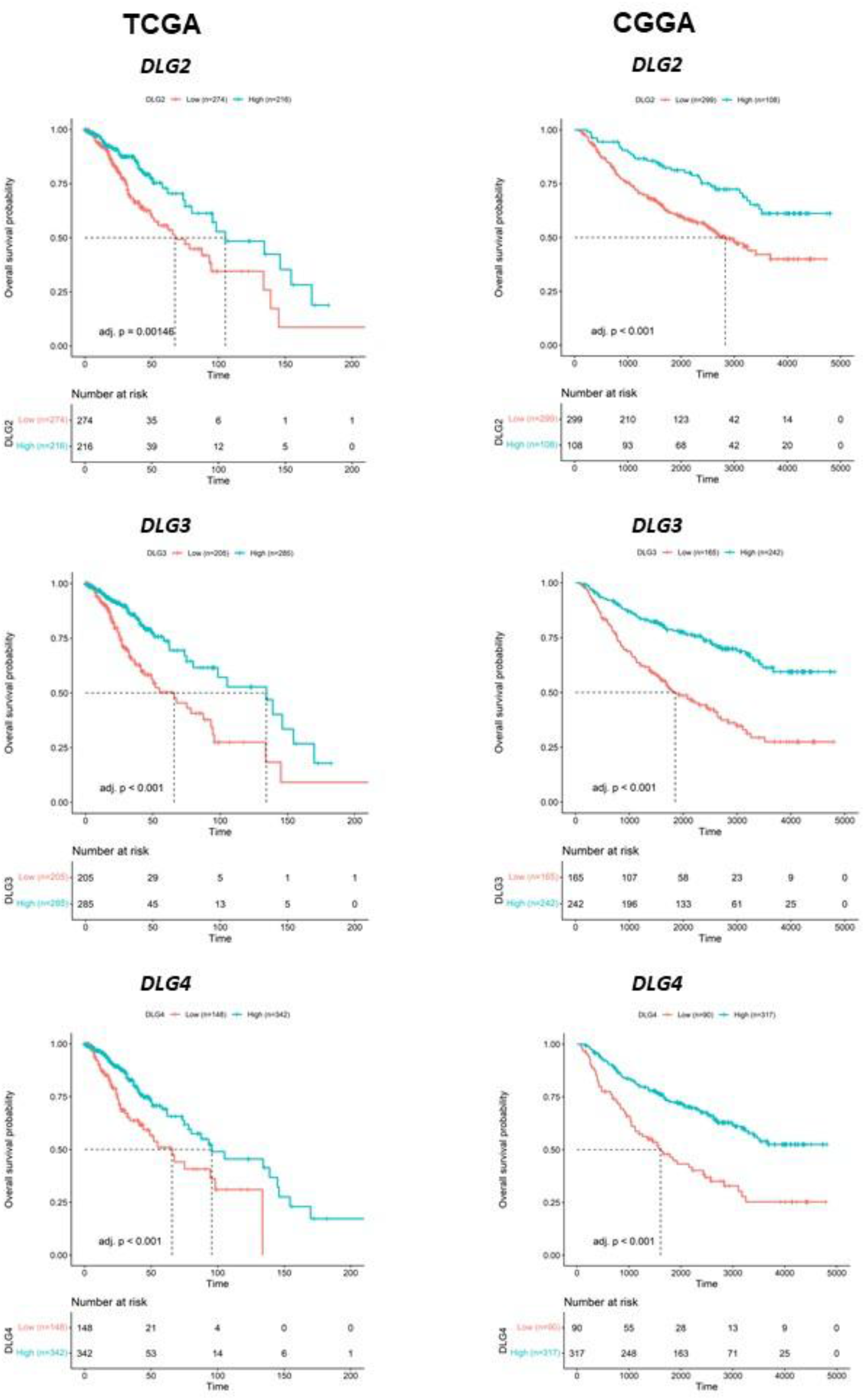
Kaplan-Meier analysis of OS in patients bearing TCGA (left) or CGGA (right) LGG tumors with higher or lower expression of *DLG2, DLG3*, and *DLG4*. The number of samples and adjusted *p*-values are indicated in the panels.

### 3.3 Expression of DLG2-DLG4 is Positively Correlated with Expression of Genes Associated with Excitatory Synaptic Transmission and Synaptic Plasticity

Among the genes included in our synaptic panel, moderate to strong positive correlations were found between 21 non-*DLG* genes and *DLG2* and between 24 non-*DLG* genes and either *DLG3* or *DLG4*. Genes showing correlations with all three *DLG* genes were *CAMK2A, CAMK2B, DLGAP1, GRIN1, GRIN2B, HOMER1, NEFL, NEFM, PPP3CA, PRKCG, RBFOX3, SHANK1, SHANK2, SHANK3, SLC17A6, SLC17A7, SYNGAP1*, and *TBR1*. There were also positive correlations in expression among the *DLG* genes themselves (Supplementary Figure S7, Table 1). Moreover, each individual *DLG* gene showed a positive correlation with the overall synaptic gene program derived from the full gene set (Figure 4).

**Table 1.**
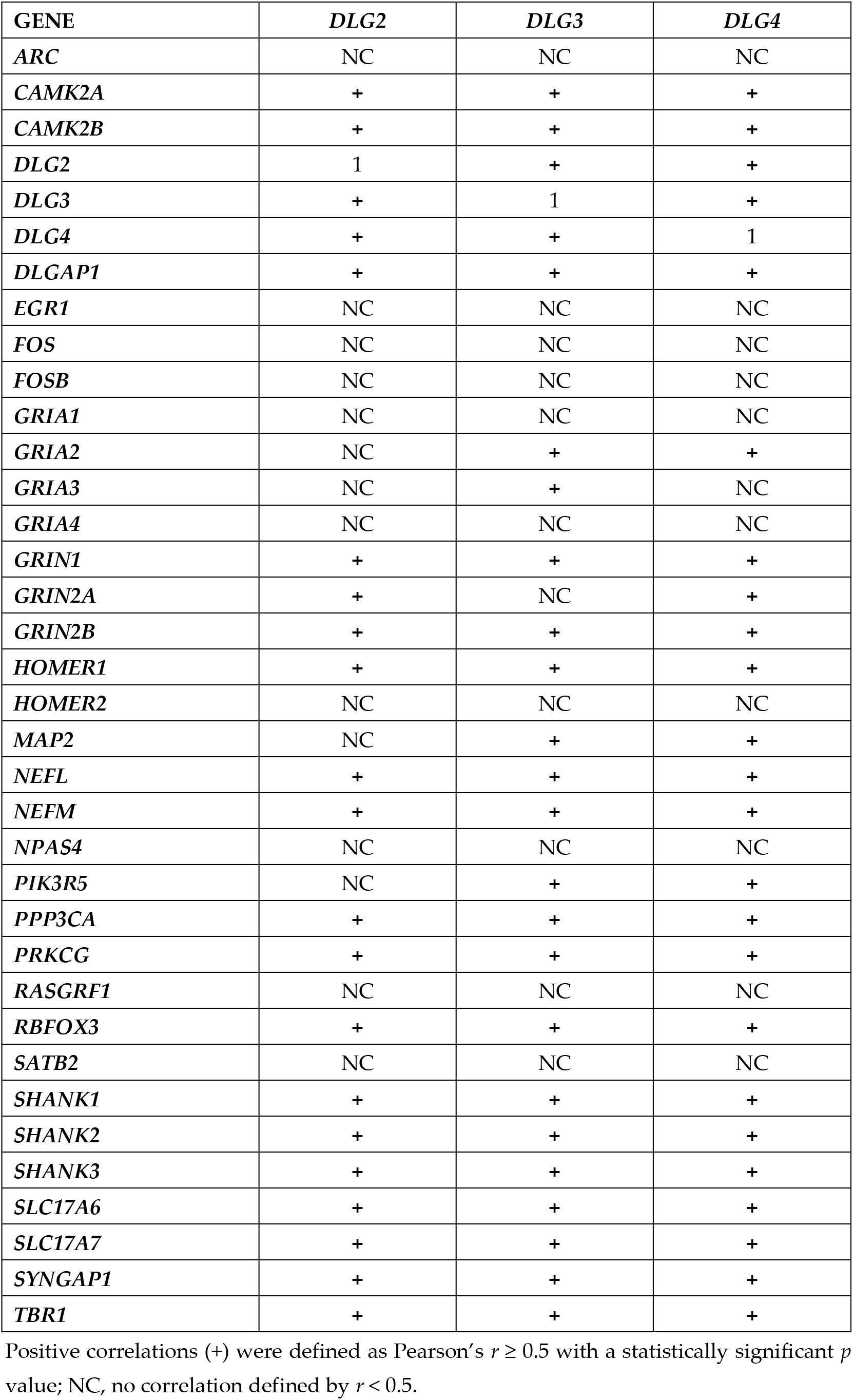
Correlations between expression of *DLG2*–*DLG4* and genes involved in excitatory synaptic transmission and synaptic plasticity.

| GENE | *DLG2* | *DLG3* | *DLG4* |
| --- | --- | --- | --- |
| *ARC* | NC | NC | NC |
| *CAMK2A* | + | + | + |
| *CAMK2B* | + | + | + |
| *DLG2* | 1 | + | + |
| *DLG3* | + | 1 | + |
| *DLG4* | + | + | 1 |
| *DLGAP1* | + | + | + |
| *EGR1* | NC | NC | NC |
| *FOS* | NC | NC | NC |
| *FOSB* | NC | NC | NC |
| *GRIA1* | NC | NC | NC |
| *GRIA2* | NC | + | + |
| *GRIA3* | NC | + | NC |
| *GRIA4* | NC | NC | NC |
| *GRIN1* | + | + | + |
| *GRIN2A* | + | NC | + |
| *GRIN2B* | + | + | + |
| *HOMER1* | + | + | + |
| *HOMER2* | NC | NC | NC |
| *MAP2* | NC | + | + |
| *NEFL* | + | + | + |
| *NEFM* | + | + | + |
| *NPAS4* | NC | NC | NC |
| *PIK3R5* | NC | + | + |
| *PPP3CA* | + | + | + |
| *PRKCG* | + | + | + |
| *RASGRF1* | NC | NC | NC |
| *RBFOX3* | + | + | + |
| *SATB2* | NC | NC | NC |
| *SHANK1* | + | + | + |
| *SHANK2* | + | + | + |
| *SHANK3* | + | + | + |
| *SLC17A6* | + | + | + |
| *SLC17A7* | + | + | + |
| *SYNGAP1* | + | + | + |
| *TBR1* | + | + | + |

Positive correlations (+) were defined as Pearson's $r \geq 0.5$ with a statistically significant $p$ value; NC, no correlation defined by $r < 0.5$ .

**Figure 4.**
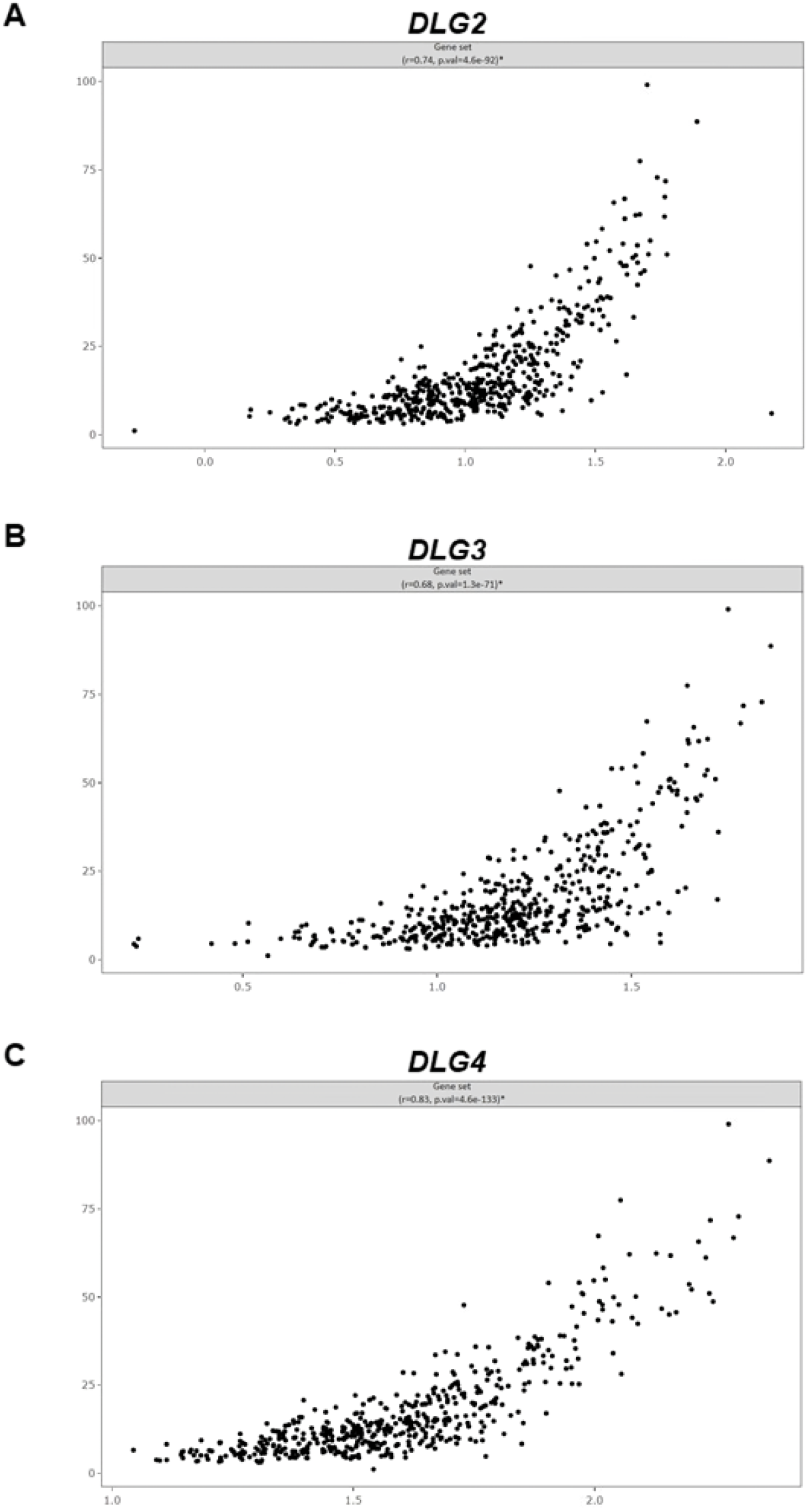
Expression of (**A**) *DLG2*, (**B**) *DLG3*, and (**C**) *DLG4* correlates with a postsynaptic excitatory transmission and synaptic plasticity gene signature, represented by a composite gene set score obtained from all genes in the panel, in TCGA LGG tumors. The horizontal axis represents *DLG* gene expression and the vertical axis represents the gene set score. Pearson’s *r* and *p*-values are shown in the panels.

A heatmap showing expression correlations of a synaptic gene set in TCGA LGG tumors indicated a non-random co-expression structure across samples. Hierarchical clustering of Spearman correlation coefficients revealed a modular organization of gene expression. Distinct clusters corresponded to key functional synaptic components, including postsynaptic scaffolding proteins, which include *DLG2-DLG4* and *SHANK* family genes, glutamatergic receptor subunits such as *GRIA* and *GRIN* genes, and activity-dependent transcriptional regulators such as *ARC, EGR1*, and *FOS*. These patterns indicate coordinated regulation of synaptic gene programs within glioma transcriptomes (Figure 5).

**Figure 5.**
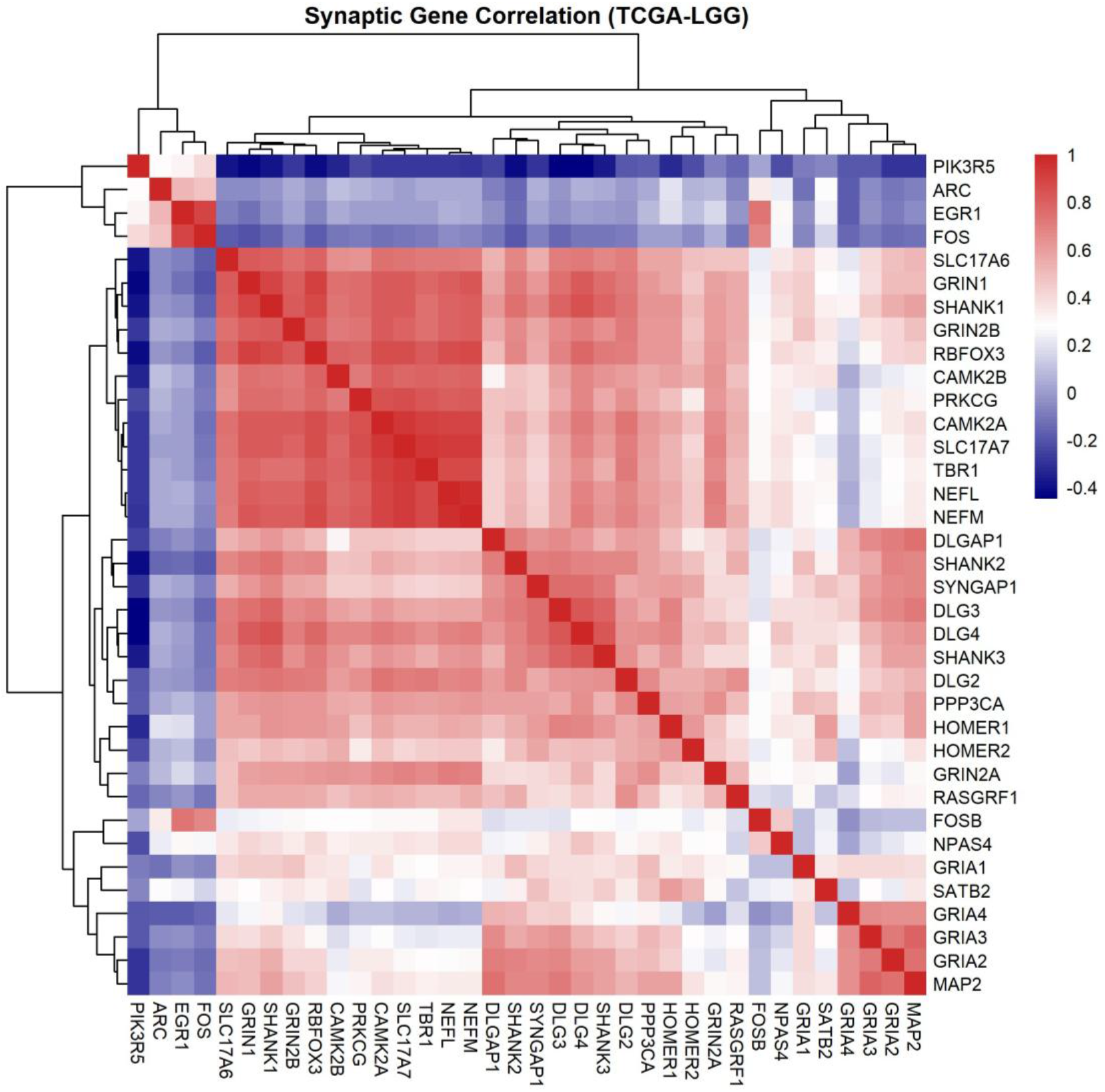
Co-expression structure of *DLG2-DLG4* and other synaptic genes in glioma. Pairwise correlations of the curated synaptic gene set were computed in TCGA LGG using RNA-seq data processed via Genomic Data Commons (GDC)/TCGAbiolinks. Expression values were log2-transformed and z-scored across samples. Spearman correlation coefficients were calculated for all gene pairs to generate a gene–gene correlation matrix. Genes were hierarchically clustered using Ward’s method and visualized as a heatmap (pheatmap, R). Color scale reflects correlation strength (blue, negative; red, positive).

## 4. Discussion

LGGs, especially IDH-mutant astrocytomas, can eventually progress to aggressive, malignant tumors with GBM-like features [4, 13, 21]. Increasing attention has been given to neural transmission and plasticity pathways co-opted by GBM tumors to stimulate brain growth [1-3]. In contrast, synaptic components in LGG remain to be characterized. We provide early evidence, based on mRNA transcription as a measure of gene expression, suggesting that excitatory neurotransmission and post-synaptic plasticity features may be present in LGG tumors and associate with patient prognosis.

*DLG2, DLG3*, and *DLG4* encode the MAGUK scaffold proteins PSD-93, SAP-102, and PSD-95. These are core proteins of the PSD at excitatory synapses, where they serve as organizational hubs that assemble and stabilize signaling complexes. All three proteins share a modular structure (PDZ, SH3, and GK domains) that allows them to bind multiple partners simultaneously. Through these interactions, they anchor and cluster NMDA and AMPA receptors at the synaptic membrane and link them to intracellular signaling pathways. They also connect receptors to other scaffold proteins such as Shank1-3, helping maintain synapse structure and efficiency. In excitatory synaptic transmission, MAGUK proteins ensure proper receptor localization and density at the postsynaptic membrane, which directly influences synaptic strength and responsiveness to glutamate release. In synaptic plasticity, especially LTP, these proteins play a central role by regulating activity-dependent changes in receptor trafficking and signaling. PSD-95 (*DLG4*) is particularly important for stabilizing AMPA receptors at synapses and promoting their insertion during LTP, whereas PSD-93 (*DLG2*) participates in organizing NMDA receptor–associated signaling complexes and can modulate synaptic strength, partly through interactions with PSD-95. SAP-102 (*DLG3*) is more prominent during development, supporting early synapse formation and plasticity by regulating NMDA receptor trafficking [7-12, 22, 23]. The role of MAGUK proteins in synaptic function leads to implications in memory formation and neurodegenerative diseases [24, 25].

We found consistent associations of all three *DLG* genes with better prognosis, as assessed by longer OS in patients with LGG, which is somewhat counterintuitive given the proposed contributing role of synaptic mechanisms in glioma progression indicated by studies in GBM. If there is any causal relationship between high *DLG2-DLG4* expression and survival, which cannot be concluded based on our current results, it is possible that *DLG* genes and MAGUK proteins, together with their associated network of synaptic components, confer a more differentiated, neuron-like transcription profile to LGG tumors, thus reducing their aggressiveness and likelihood of progressing towards GBM. In neuroblastoma, it has been shown that low *DLG2* expression induces forced cell cycle progression and predicts poor patient survival [26]. Differentiation of cells representative of different glioma types, including grade 3 astrocytoma and oligodendroglioma, into neuron-like cells, can be induced by a combination of small molecules or overexpression of neural transcription factors. This promotion of neuronal differentiation reduces cell proliferation and expression of tumor-associated proteins, while upregulating tumor suppression genes [27]. Other evidence supports the view that enrichment of neuronal differentiation-related genes may be a therapeutic strategy to suppress the malignant phenotype and improve prognosis in glioma [28]. In addition, our correlation analyses indicate a modular and non-random co-expression architecture between *DLG*-*DLG4* and other synaptic genes, with co-ordinated clusters in the gene set corresponding to synaptic scaffolding, glutamatergic signaling, and activity-dependent transcriptional programs. It is noteworthy that, despite tumors with higher expression of *DLG2-DLG4* have better prognosis, the gene expression levels still do not seem to reach those of non-tumor brain tissue.

LGGs and GBM tumors are biologically related neoplasms that share core features of diffuse infiltration, genetic instability, and adaptation within the brain microenvironment. GBM has been conceptualized as the terminal stage of malignant progression from LGGs, particularly in tumors harboring mutations in *IDH1* or *TP53*. Integrated molecular profiling has demonstrated that GBMs can arise as primary tumors biologically distinct from IDH-mutant LGGs. IDH-mutant LGGs generally display slower growth kinetics, a hypermethylated epigenetic landscape, and more prolonged evolutionary trajectories. Thus, LGGs and glioblastomas may exist along partially overlapping but biologically divergent evolutionary pathways. These distinctions may help explain why differentiation toward a neuronal-like phenotype might be associated with reduced aggressiveness in LGGs (Figure 6), whereas similar transcriptional programs have been linked to increased malignancy and cellular plasticity in GBM [3, 13, 17].

**Figure 6.**
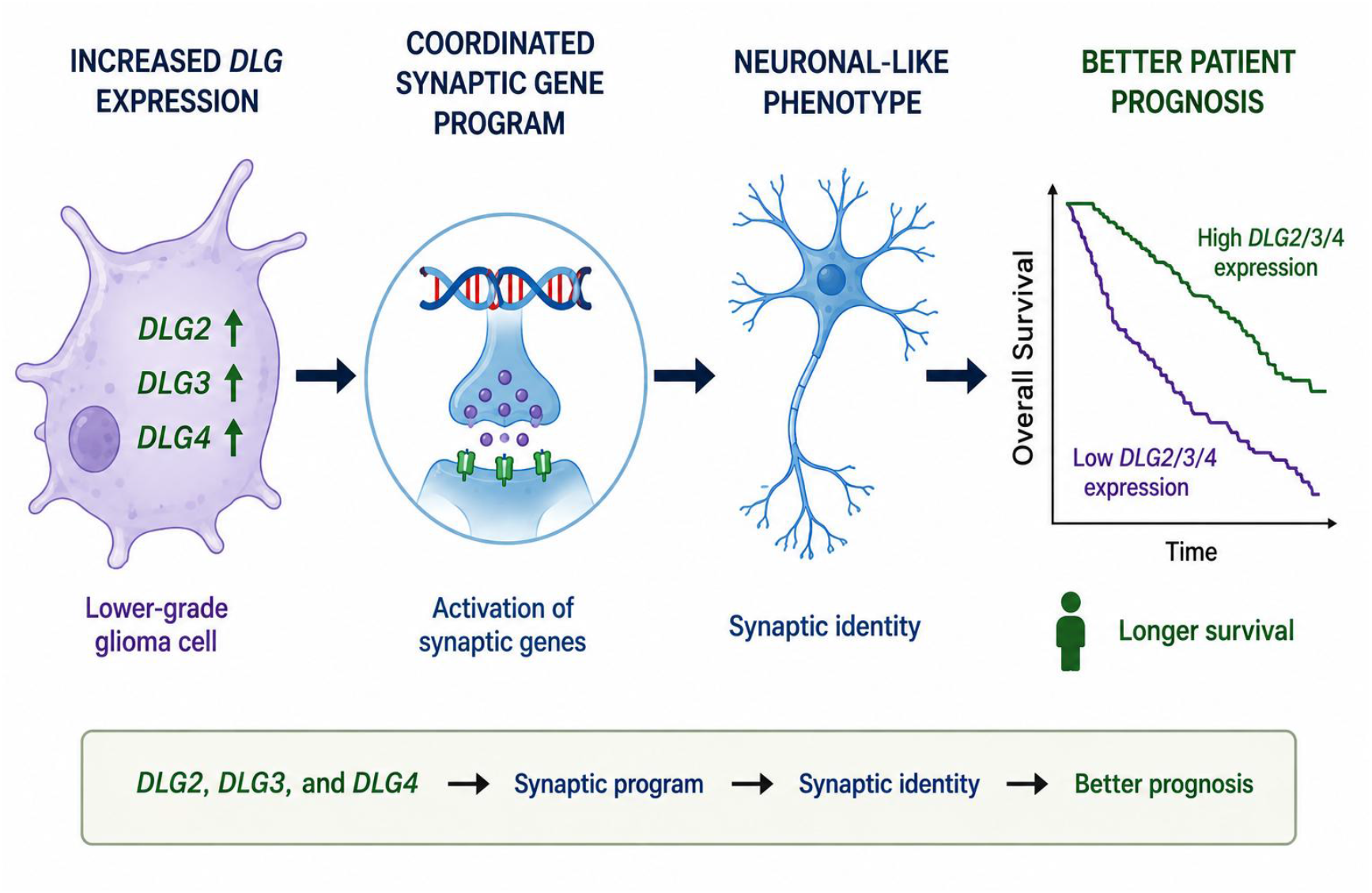
Proposed hypothetical model linking increased expression of *DLG2, DLG3*, and *DLG4* with synaptic-associated transcriptional programs and favorable prognosis in LGG.

Our study has several limitations. First, for some of the analyses we used the glioma histological classification, which is now outdated. The definitions for LGG entities were updated in the WHO 2021 classification to integrate molecular signatures with histological features [17, 29]. The histological classification of gliomas in the TCGA LGG dataset is based on WHO 2007/2016 criteria and therefore does not fully reflect the integrated molecular classification introduced in the WHO 2021 guidelines. This limitation is inherent to TCGA and other datasets, which were generated prior to the adoption of the 2021 classification. However, retaining the original histological annotations ensures reproducibility and consistency with prior TCGA-based studies, and these annotations remain relevant as they represent well-characterized morphological entities that form the basis of much of the existing glioma literature. The WHO 2021 classification recognizes that legacy datasets generated before its implementation may not be fully compatible with the updated integrated framework and explicitly supports the continued use of prior WHO histological classifications in such contexts [17]. In addition, we have also included analyses using updated classifications that incorporate molecular features. Specific mutations can influence LGG prognosis [30-33]. For example, IDH status separates LGG tumors into biologically and clinically distinct diseases, with IDH-mutated tumors having more favorable prognosis [13]. We found that, at least in the CGGA cohort, *DLG2-DLG4* genes retained associations with favorable prognosis after adjustment for IDH mutation status, 1p/19q codeletion status, age, and tumor grade. Molecular subtype analyses were performed independently according to IDH mutation and 1p/19q co-deletion status, consistent with the WHO 2021 integrated classification framework. LGG-IDH-mut-codel tumors correspond to oligodendrogliomas, whereas LGG-IDH-mut-non-codel tumors correspond to astrocytomas. Tumors classified as LGG-IDH-wt were retained as a separate category because TCGA and CGGA legacy datasets include lower-grade diffuse gliomas lacking IDH mutations, although many of these tumors would currently be considered biologically closer to GBM under contemporary molecular criteria [29].

Another limitation is that associations between gene expression and patient prognosis, or correlations with expression of other genes, do not imply mechanistic roles and causative effects. These associations should be interpreted primarily as a basis for generating mechanistic hypotheses and identifying candidate prognostic factors. Experiments examining the effects of silencing *DLG2-DLG4* in cultured LGG cells are required to demonstrate a role for these genes in reducing LGG aggressiveness. Also, it is well established that higher gene expression at the mRNA level does not necessarily represent increased protein content. Some studies specifically addressing this issue in different model systems have reported broad agreement between mRNA and protein levels, whereas others have found a weak positive correlation [34-36]. Further research is required to examine the expression and possible role of *DLG2-DLG4*-encoded proteins in LGG.

## 5. Conclusions

High expression of *DLG2-DLG4* shows a consistent association with better OS in patients with LGG, suggesting the possibility of a prognostic role. In addition, *DLG2-DLG4* gene expression is correlated with a signature of components of neural transmission and synaptic plasticity, raising the possibility that LGG tumors may replicate cellular mechanisms involved in normal neuronal function, and features of neuronal differentiation may be related to better prognosis.

## Supporting information

Supplementary Information-Gaia et al

## Supplementary Materials

The following supporting information can be downloaded: **Supplementary Table S1:** Panel of genes chosen as representative of an excitatory synapse and synaptic plasticity signature; **Supplementary Table S2**. Cut-off values for high and low gene expression levels used for survival analyses. **Supplementary Figure S1:** Gene expression levels of (**A**) *DLG2*, (**B**) *DLG3*, and (**C**) *DLG4G* in glioma types and non-tumor brain tissue; **Supplementary Figure S2:** Gene expression levels of (**A**) *DLG2*, (**B**) *DLG3*, and (**C**) *DLG4* in CGGA LGG tumors classified into histological types; **Supplementary Figure S3:** Gene expression levels of (**A**) *DLG2*, (**B**) *DLG3*, and (**C**) *DLG4* in TCGA LGG tumors classified into molecular subtypes; **Supplementary Figure S4**.

Expression levels of *DLG2, DLG3*, and *DLG4* in relation to PC1 in TCGA LGG tumors; Supplementary Figure S5. Forest plots showing multivariable Cox proportional hazards analyses for OS in TCGA LLG tumors; **Supplementary Figure S6**. Forest plots showing multivariable Cox proportional hazards analyses for OS in CGGA LLG tumors; **Supplementary Figure S7:** Correlations found between expression of (**A**) *DLG2*, (**B**) *DLG3*, and (**C**) *DLG4* and individual genes from the synaptic dataset in TCGA LGG tumors; **Supplementary references**.

## Author Contributions

Conceptualization, F.G., R.R. and G.I.R.; methodology, F.G., H.R.D.P., R.R. and G.R.I; formal analysis, F.G., H.R.D.P. and R.R.; investigation, F.G., H.R.D.P., R.R. and G.R.I.; resources, F.G., O.M., R.R. and G.R.I.; data curation, F.G., H.R.D.P. and R.R.; writing—original draft preparation, H.R.D.P. and R.R.; writing—review and editing, F.G., H.R.D.P., O.M., R.R. and G.R.I.; supervision, R.R. and G.R.I.; project administration, R.R. and G.R.I.; funding acquisition, F.G., O.M., R.R. and G.R.I. All authors have read and agreed to the published version of the manuscript.

## Funding

This work was supported by the National Council for Scientific and Technological Development (CNPq, MCTI, Brazil) grant numbers 304623/2025-3 and 406484/2022–8 (INCT BioOncoPed) to R.R., the Children’s Cancer Institute (ICI), Brazilian Federal Agency for Support and Evaluation of Graduate Education (CAPES), The Center for Advanced Neurology and Neurosurgery (CEANNE), and the Mackenzie Evangelical University.

## Institutional Review Board Statement

Not applicable.

## Informed Consent Statement

Not applicable.

## Data Availability Statement

This research used data generated by The Cancer Genome Atlas (TCGA) Research Network (https://www.cancer.gov/tcga) and the Chinese Glioma Genome Atlas (CGGA, http://www.cgga.org.cn), including TCGA data accessed through the Evergene platform (https://bshihlab.shinyapps.io/evergene/). Data from the Rembrandt dataset were also used through the GlioVis platform (https://gliovis.bioinfo.cnio.es/).

## Conflicts of Interest

The authors declare no conflicts of interest related to the contents of this study. The funders had no role in the design of the study; in the collection, analyses, or interpretation of data; in the writing of the manuscript; or in the decision to publish the results.

## Abbreviations

The following abbreviations are used in this manuscript:

AMPA: α-amino-3-hydroxy-5-methyl-4-isoxazolepropionic acid
CGGA: Chinese Glioma Genome Atlas
Codel: Co-deletion
GBM: Glioblastoma
G-CIM: Glioma CpG island methylator phenotype
GDC: Genomic Data Commons
HR: Hazard Ratio
LGG: Lower-grade glioma
LGG-IDH-mut-codel: Lower-grade glioma with a mutation in the isocitrate dehydro-genase (IDH) gene and co-deletion of chromosomal arms 1p and 19q
LGG-IDH-mut-non-codel: Lower-grade glioma with a mutation in the IDH gene without co-deletion of chromosomal arms 1p and 19q
LGG-IDH-wt: Lower-grade glioma wild-type for IDH
LTP: Long-term potentiation
MAGUK: Membrane-associated guanylate kinase
NA: Not available
NC: No correlation
NMDA: N-methyl-D-aspartate
OS: Overall survival
PA: Pilocytic astrocytoma
PC1: First principal component
PCA: Principal Component Analysis
TCGA: The Cancer Genome Atlas
WHO: World Health Organization

## Notes

https://www.cancer.gov/tcga

http://www.cgga.org.cn

https://bshihlab.shinyapps.io/evergene/

https://gliovis.bioinfo.cnio.es/

### Competing Interest Statement

The authors have declared no competing interest.

### Summary of Updates

The manuscript has been extensively extended, revised and improved. New analyses were included both in the main and supplementary information files.

