## Supplementary Information-Gaia et al for "*DLG2–DLG4* Expression is Associated with Improved Survival and a Synaptic Gene Signature in Lower-Grade Glioma"

**Supplementary Table S1.** Panel of genes chosen as representative of an excitatory synapse and synaptic plasticity signature.

| GENE | PROTEIN | SYNAPTIC FUNCTIONS | REFERENCES |
|---|---|---|---|
| ***ARC*** | Activity-regulated cytoskeleton-associated protein | AMPAR endocytosis, synaptic scaling, plasticity, long-term memory | [S1, S2] |
| ***CAMK2A*** | CaMKII alpha | LTP induction, AMPAR phosphorylation | [S3, S4] |
| ***CAMK2B*** | CaMKII beta | Actin binding, spine structural plasticity, LTP | [S5, S6] |
| ***DLG2*** | PSD-93 (SAP-93) | Scaffolds excitatory receptors at synapses | [5, 6, S7, S8] |
| ***DLG3*** | SAP-102 | Regulates NMDA receptor trafficking, maturation | [4, S8, S9] |
| ***DLG4*** | PSD-95 | Anchors receptors, drives synaptic plasticity | [8, S8, S10] |
| ***DLGAP1*** | SAPAP1 (GKAP) | PSD scaffold linking PSD95–SHANK | [S11, S12] |
| ***EGR1*** | Early growth response protein 1 | Activity-dependent transcription, plasticity | [S13, S14] |
| ***FOS*** | c-Fos | Immediate early gene, activity transcription | [S15] |
| ***FOSB*** | FosB proto-oncogene | Sustained activity-dependent transcription factor | [S16, S17] |
| ***GRIA1*** | GluA1 (GluR1) | AMPAR subunit, synaptic strengthening | [S18, S19] |
| ***GRIA2*** | GluA2 (GluR2) | AMPAR $Ca^{2+}$ impermeability, channel regulation | [S20, S21] |
| ***GRIA3*** | GluA3 (GluR3) | AMPAR subunit, basal transmission | [S22, S23] |
| ***GRIA4*** | GluA4 (GluR4) | AMPAR subunit, fast excitatory transmission | [S22, S24] |
| ***GRIN1*** | GluN1 | Essential NMDAR subunit, channel function | [S25, S26] |
| ***GRIN2A*** | GluN2A | NMDAR subunit, synaptic maturation | [S27, S28] |
| ***GRIN2B*** | GluN2B | NMDAR subunit, plasticity signaling | [S27, S29] |
| ***HOMER1*** | Homer1 | Links mGluRs to PSD scaffold | [S29, S30] |
| ***HOMER2*** | Homer2 | Scaffold for mGluR signaling complexes | [S31, S32] |
| ***MAP2*** | Microtubule-associated protein 2 | Dendritic stability, microtubule organization | [S33, S34] |
| ***NEFL*** | Neurofilament light polypeptide | Axonal structure, cytoskeletal support | [S35, S36] |
| ***NEFM*** | Neurofilament medium chain | Axonal caliber, structural integrity | [S37] |
| ***NPAS4*** | Neuronal PAS domain protein 4 | Activity-dependent synaptic gene regulation | [S38, S39] |

| | | | |
|---|---|---|---|
| ***PIK3R5*** | PI3K regulatory subunit gamma | PI3K signaling, synaptic plasticity | [S40, S41] |
| ***PPP3CA*** | Calcineurin A alpha | Activity-dependent phosphatase, LTD signaling | [S42, S43] |
| ***PRKCG*** | PKC gamma | Modulates synaptic signaling, plasticity | [S44, S45] |
| ***RASGRF1*** | RasGRF1 | Ras activation, synaptic plasticity signaling | [S46, S47] |
| ***RBFOX3*** | NeuN | Neuronal splicing factor, identity marker | [S48, S49] |
| ***SATB2*** | SATB2 | Cortical neuron identity, neuronal development, gene regulation | [S50, S51] |
| ***SHANK1*** | Shank1 | PSD scaffold, spine structure | [S52, S53] |
| ***SHANK2*** | Shank2 | PSD scaffold, receptor organization | [S53, S54] |
| ***SHANK3*** | Shank3 | PSD scaffold, synaptic stability | [S53, S55] |
| ***SLC17A6*** | VGLUT2 | Vesicular glutamate transporter | [S56, S57] |
| ***SLC17A7*** | VGLUT1 | Vesicular glutamate transporter | [S58, S59] |
| ***SYNGAP1*** | SynGAP | Ras regulation, AMPAR trafficking | [S60, S61] |
| ***TBR1*** | T-box brain protein 1 | Cortical neuron identity, glutamatergic differentiation | [S62] |

**Supplementary Table S2.** Cut-off values for high and low gene expression levels used for survival analyses.

| **Dataset** | **Gene** | **Cutpoint** | **Low expression (*n*)** | **High expression (*n*)** |
|---|---|---|---|---|
| TCGA-LGG | *DLG2* | 0.1234 | 274 | 216 |
| TCGA-LGG | *DLG3* | -0.2252 | 205 | 285 |
| TCGA-LGG | *DLG4* | -0.5811 | 148 | 342 |
| CGGA LGG (merged 325+693) | *DLG2* | 0.645 | 299 | 108 |
| CGGA LGG (merged 325+693) | *DLG3* | -0.2327 | 165 | 242 |
| CGGA LGG (merged 325+693) | *DLG4* | -0.6859 | 90 | 317 |

All expression values were standardized using gene-wise z-score transformation.

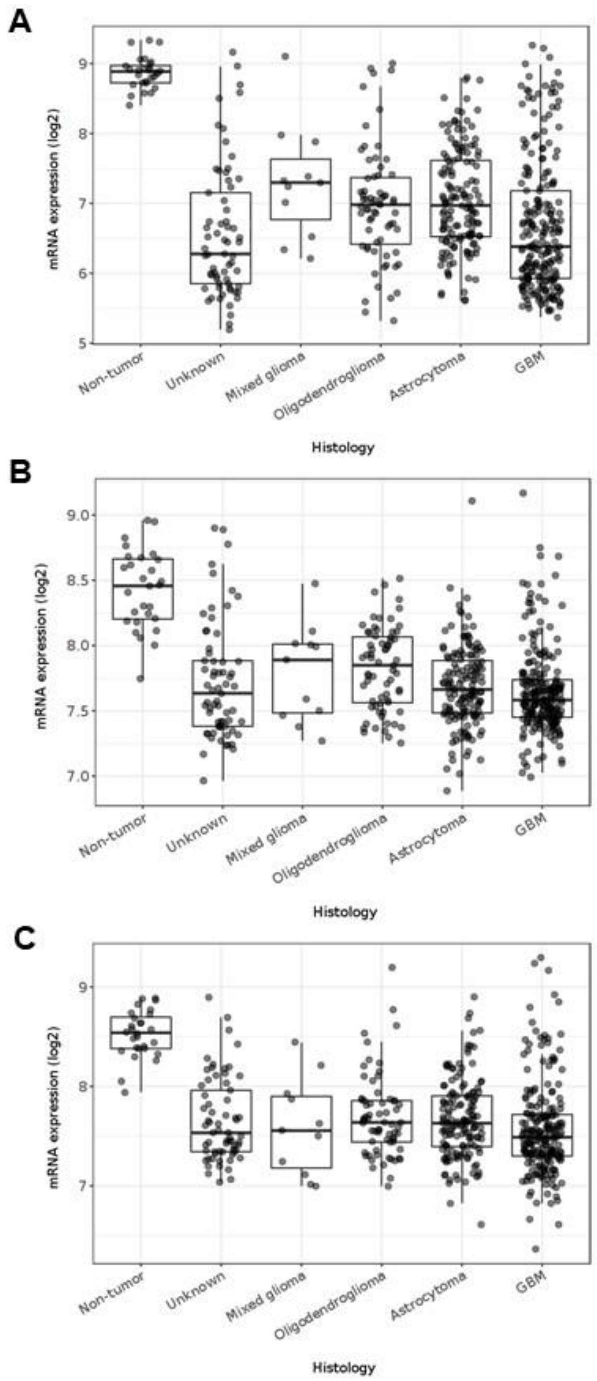


**Supplementary Figure S1.** Gene expression levels of (**A**) *DLG2*, (**B**) *DLG3*, and (**C**) *DLG4* in glioma types and non-tumor brain tissue. Data were obtained from the Rembrandt dataset [15] was analyzed using the GlioVis-Data Visualization Tools for Brain Tumor Datasets (https://gliovis.bioinfo.cnio.es/) platform [19].

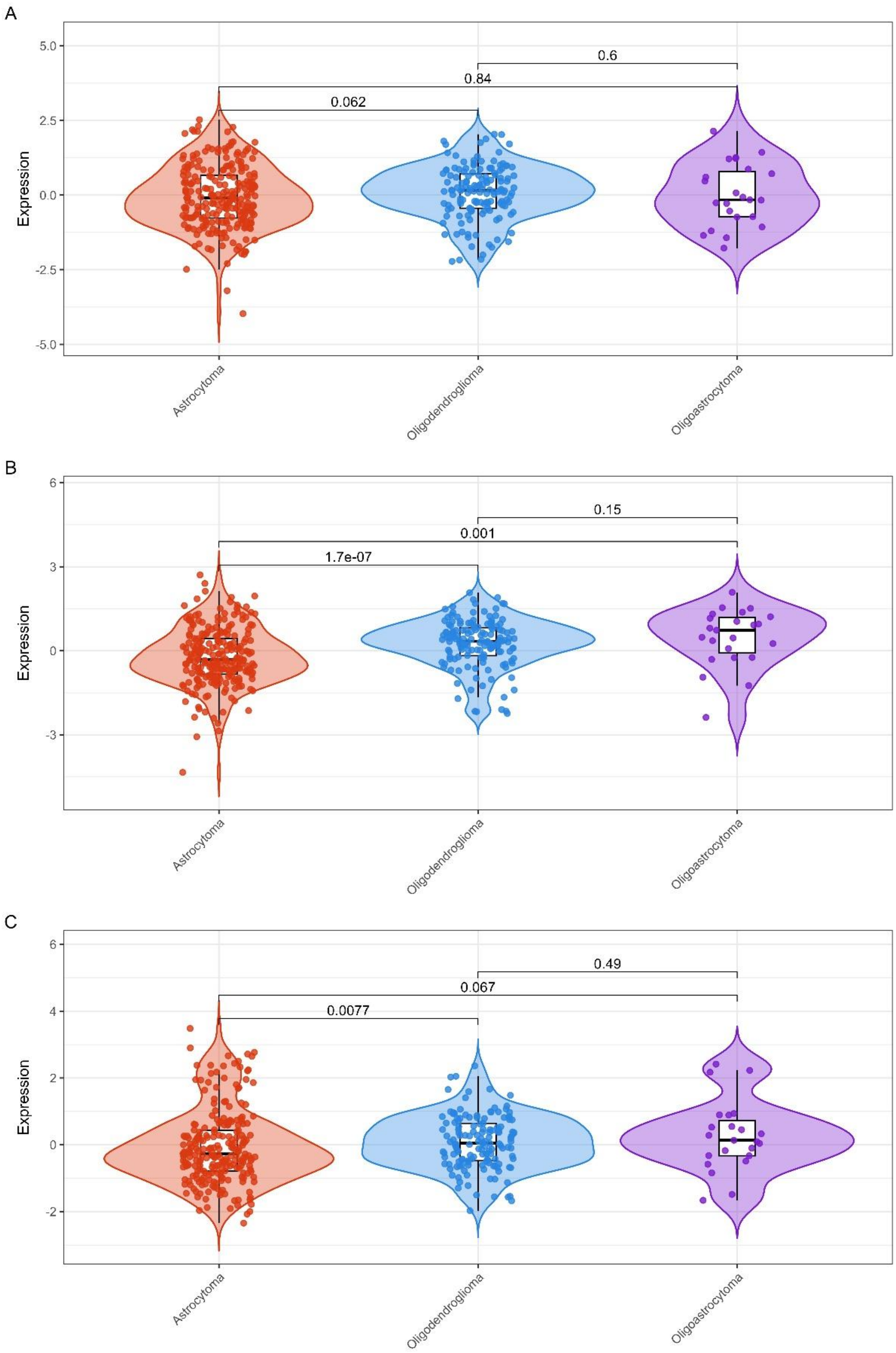


**Supplementary Figure S2.** Gene expression levels of (**A**) *DLG2*, (**B**) *DLG3*, and (**C**) *DLG4* in CGGA LGG tumors classified into histological types. Astrocytoma, n = 239; oligodendroglioma, n = 145; oligoastrocytoma, n = 23; *p* values are indicated in the panels.

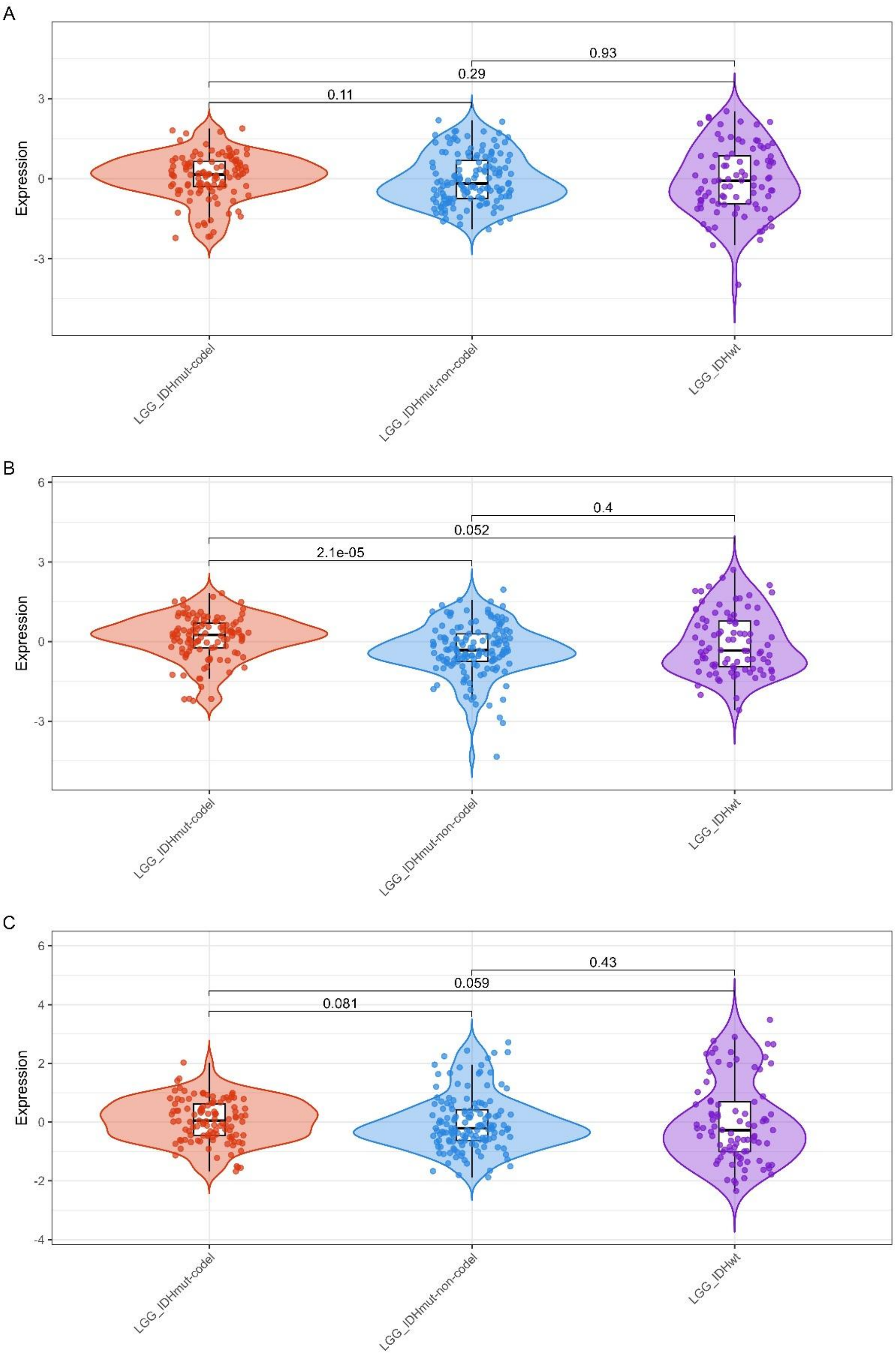


**Supplementary Figure S3.** Gene expression levels of (**A**) *DLG2*, (**B**) *DLG3*, and (**C**) *DLG4* in CGGA LGG tumors classified into molecular subtypes. LGG-IDH-mut-codel, n = 113; LGG-IDH-mut-non-codel, n = 142; LGG-IDH-wt, n = 88; $p$ values are indicated in the panels.

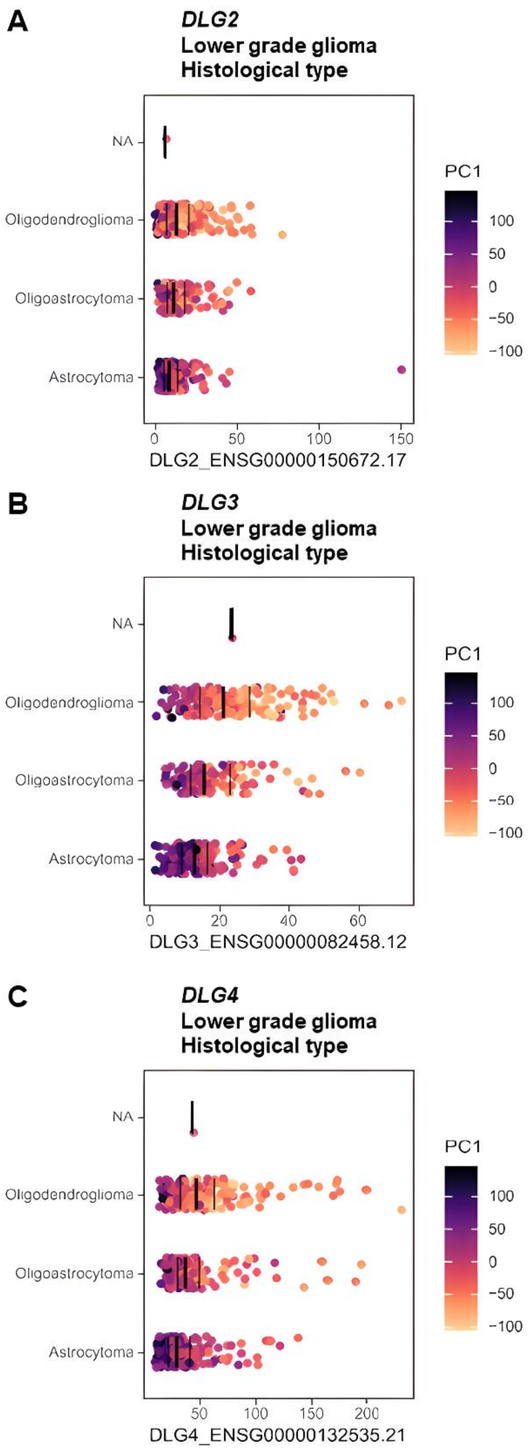
A
DLG2
Lower grade glioma
Histological type
NA
Oligodendroglioma
Oligoastrocytoma
Astrocytoma
0
50
100
150
DLG2_ENSG00000150672.17
PC1
100
50
0
−50
−100
B
DLG3
Lower grade glioma
Histological type
NA
Oligodendroglioma
Oligoastrocytoma
Astrocytoma
0
20
40
60
DLG3_ENSG00000082458.12
PC1
100
50
0
−50
−100
C
DLG4
Lower grade glioma
Histological type
NA
Oligodendroglioma
Oligoastrocytoma
Astrocytoma
50
100
150
200
DLG4_ENSG00000132535.21
PC1
100
50
0
−50
−100

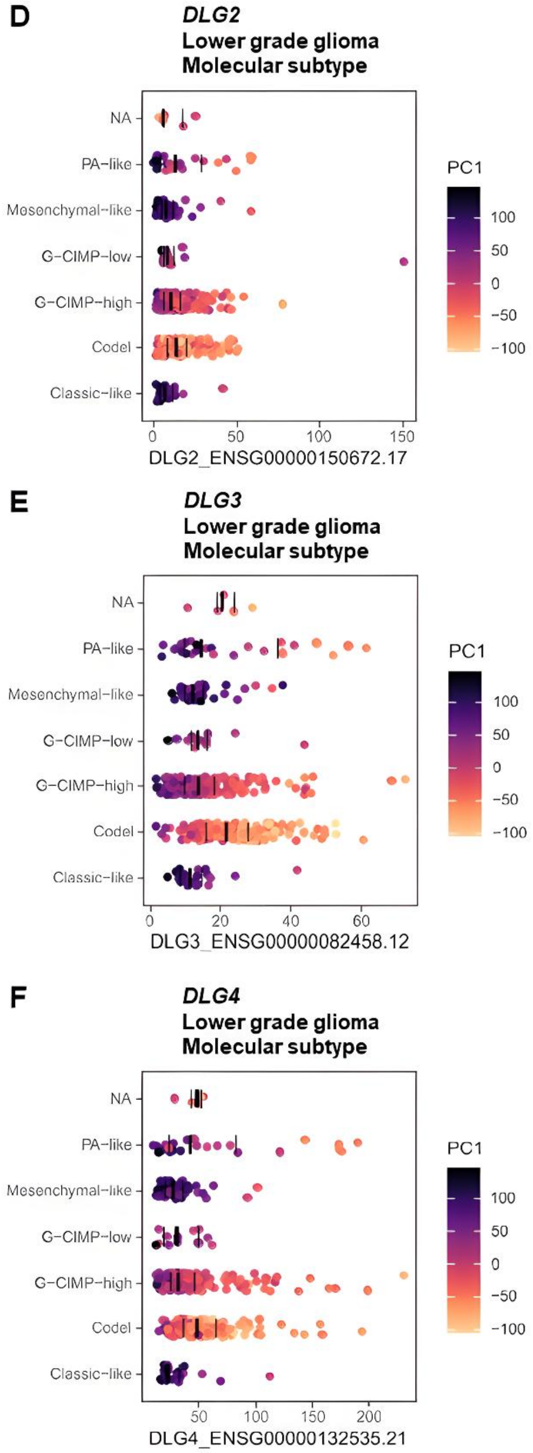


**Supplementary Figure S4.** Expression levels of *DLG2*, *DLG3*, and *DLG4* in relation to PC1 in TCGA LGG tumors. (**A**) *DLG2*, histological type; (**B**) *DLG3*, histological type; (**C**) *DLG4*, histological type; (**D**) *DLG2*, molecular subtype; (**E**) *DLG3*, molecular subtype; (**F**) *DLG4*, molecular subtype. For this analysis, molecular subtypes were classified as PA-like, mesenchymal-like, G-CIMP-high, G-CIMP-low, codel, classic-like, or NA [17, 20].

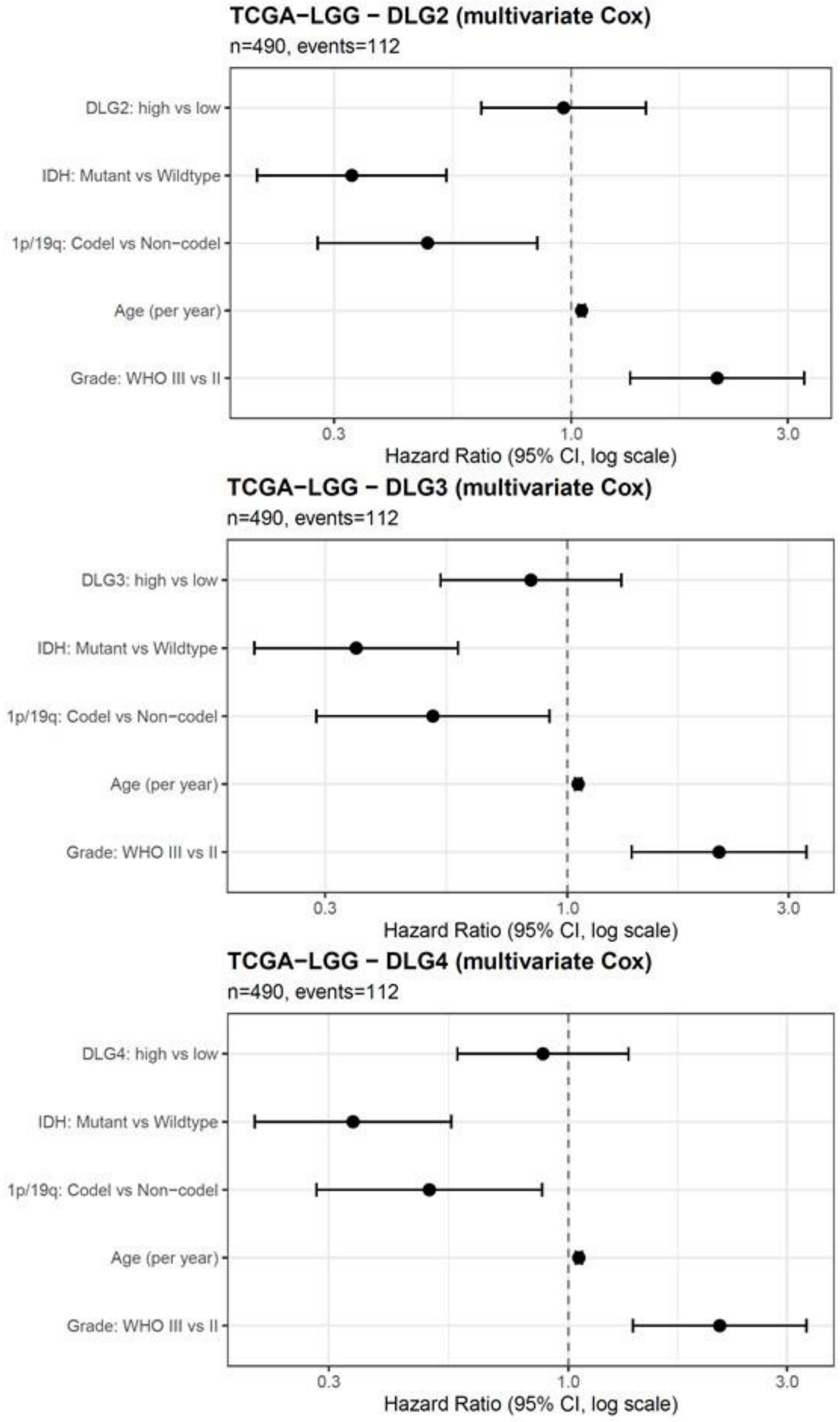


**Supplementary Figure S5.** Forest plots showing multivariable Cox proportional hazards analyses for OS in TCGA LLG tumors. Models included *DLG* gene expression status together with established clinicomolecular prognostic variables, namely IDH mutation status, 1p/19q codeletion status, age, and tumor grade. Hazard ratios (HRs) and 95% confidence intervals are shown. HR < 1 indicates favorable prognostic association, whereas HR > 1 indicates increased risk of death.

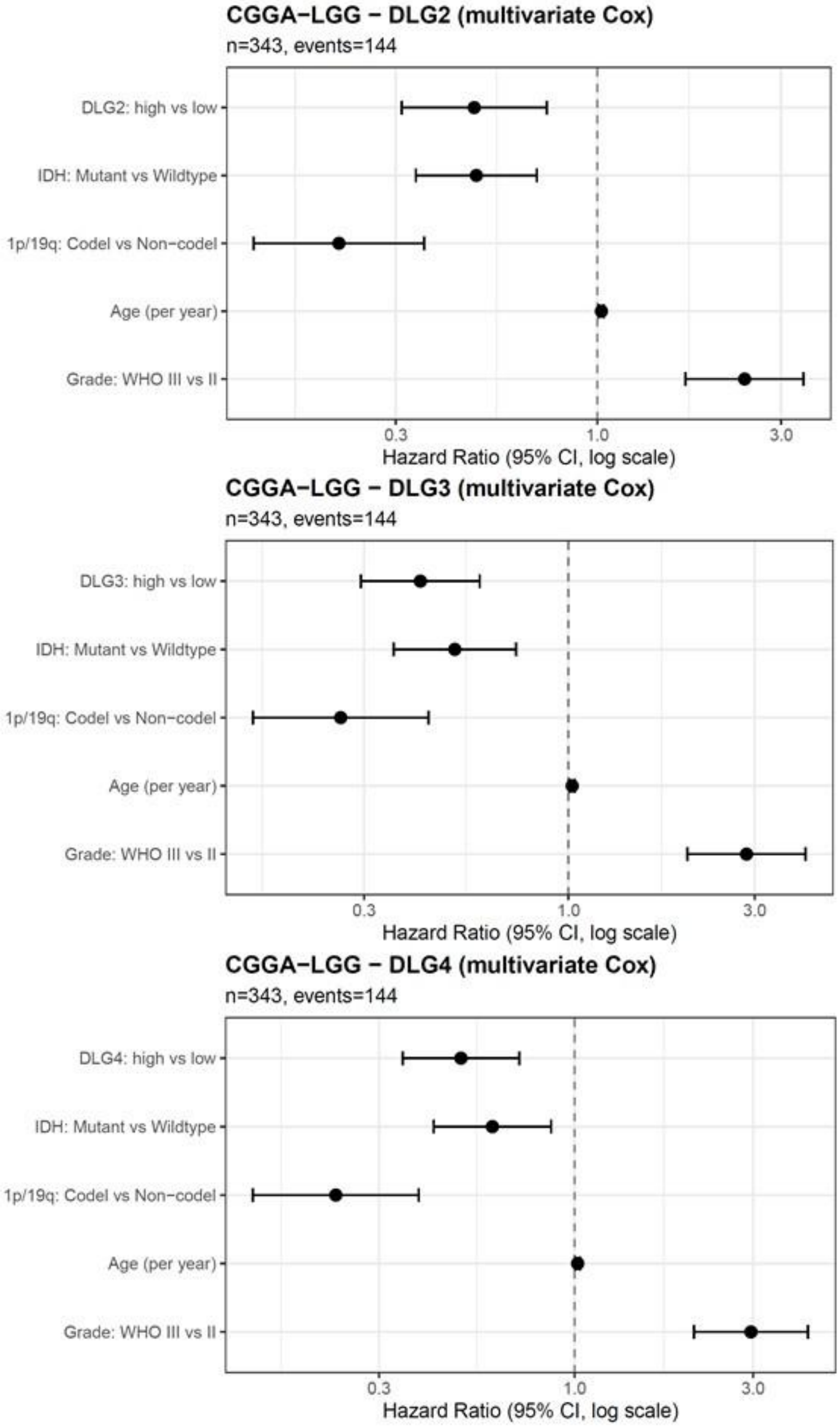


**Supplementary Figure S6.** Forest plots showing multivariable Cox proportional hazards analyses for OS in CGGA LLG tumors. Models included *DLG* gene expression status together with established clinicomolecular prognostic variables, namely IDH mutation status, 1p/19q codeletion status, age, and tumor grade. Hazard ratios (HRs) and 95% confidence intervals are shown. HR < 1 indicates favorable prognostic association, whereas HR > 1 indicates increased risk of death.

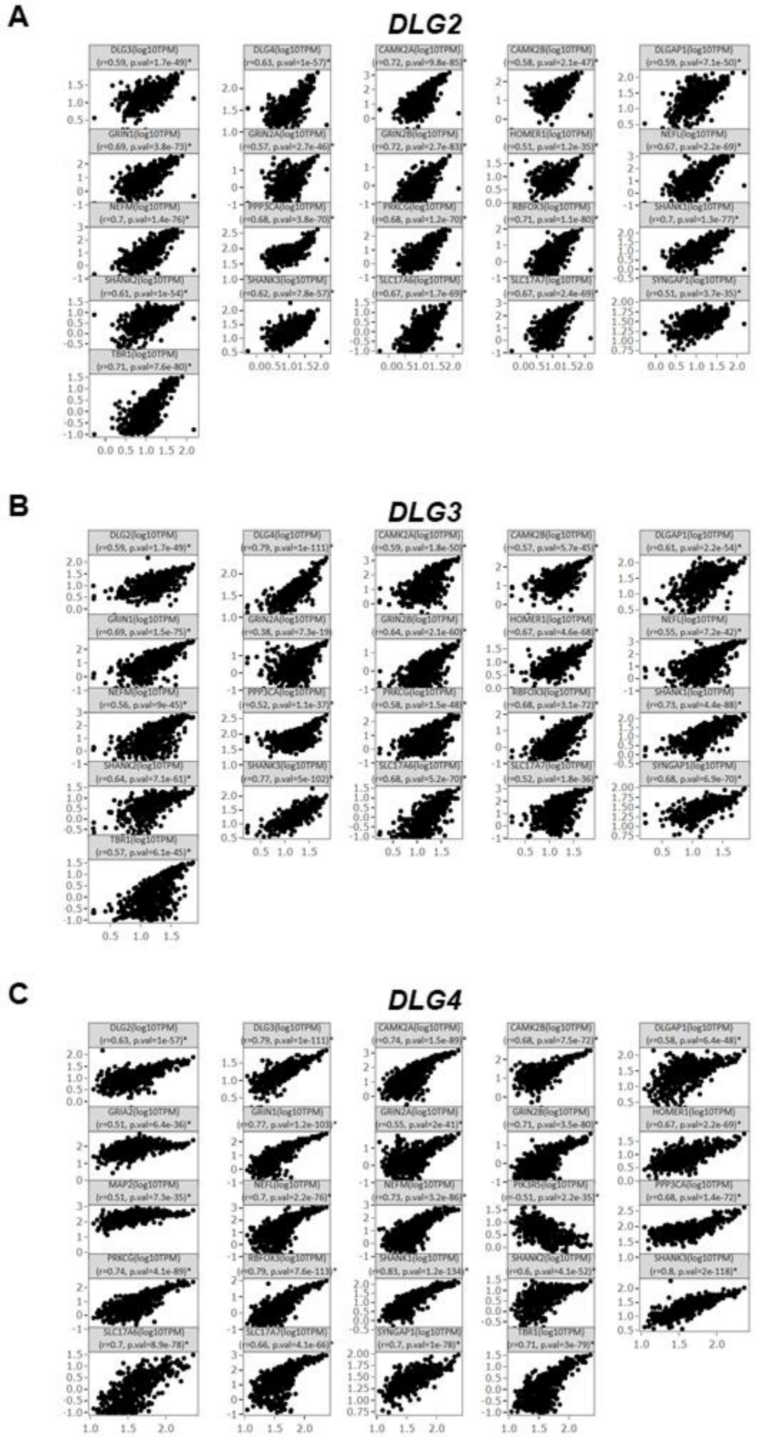


**Supplementary Figure S7.** Correlations found between expression of (**A**) *DLG2*, (**B**) *DLG3*, and (**C**) *DLG4* and individual genes from the synaptic dataset in TCGA LGG tumors; Pearson's r and *p* values are indicated in the panels.
